# Clathrin adaptor AP-1 and Stratum act in parallel pathways to control Notch activation in *Drosophila* Sensory Organ Precursor Cells

**DOI:** 10.1101/2020.04.08.033092

**Authors:** Karen Bellec, Mathieu Pinot, Isabelle Gicquel, Roland Le Borgne

## Abstract

*Drosophila* sensory organ precursors divide asymmetrically to generate pIIa/pIIb cells whose identity relies on the differential activation of Notch at cytokinesis. While Notch is present apically and basally relative to the midbody at the pIIa-pIIb interface, only the basal pool of Notch is reported to contribute to Notch activation in the pIIa cell. Correct intra-lineage signalling requires appropriate apico-basal targeting of Notch, its ligand Delta and its trafficking partner Sanpodo. We previously reported that AP-1 and Stratum regulate the intracellular trafficking of Notch and Sanpodo from the *trans*-Golgi network to the basolateral membrane. Loss of AP-1 or Stratum caused mild *Notch* gain-of-function phenotypes. Here, we report that the concomitant loss of AP-1 and Stratum results in a much more penetrant *Notch* gain-of-function phenotype indicating that AP-1 and Strat control two parallel pathways. While unequal partitioning of cell fate determinants and cell polarity were unaffected, Numb-mediated symmetry breaking is impaired. We further observed increased amounts of signaling competent Notch as well as Delta and Sanpodo at the apical pIIa-pIIb interface and the loss of the basal pool of Notch. We propose that AP-1 and Stratum operate in two parallel pathways to ensure the correct apico-basal localization of Notch controlling where receptor activation takes place.

## Introduction

Cell-cell signaling by the evolutionarily conserved Notch receptor promotes cell fate acquisition in a large variety of developmental processes in metazoans (Artavanis-Tsakonas et al., 1999; Bray, 1998; Kopan and Ilagan, 2009). In most of cases, Notch receptor is activated by transmembrane ligands present at the plasma membrane of adjacent cells. Following binding to Notch, endocytosis of the ligand induces pulling forces driving a change in the conformation of the Notch extracellular domain thereby unmasking the S2 cleavage site of Notch (Gordon et al., 2015; Langridge and Struhl, 2017; Meloty-Kapella et al., 2012; Seo et al., 2016; Shergill et al., 2012; Wang and Ha, 2013). This regulated cleavage is followed by a constitutive proteolytic cleavage of Notch by the gamma secretase complex (Mumm et al., 2000; Struhl and Adachi, 2000) giving rise to the Notch intracellular domain (NICD), a polypeptide that translocates into the nucleus to act as a transcriptional coactivator (Artavanis-Tsakonas et al., 1999; Bray, 1998; Kopan and Ilagan, 2009). As proteolytic activation of the Notch receptor is irreversible, Notch activation needs to be tightly controlled in time and in space. The model system of asymmetric cell division of the sensory organ precursors (SOPs) in the pupal notum of *Drosophila* has been instrumental to identify the site of Notch activation at the cell surface. SOPs are polarized epithelial cells that divide asymmetrically within the plane of the epithelium to generate two daughter cells whose fate depends on the differential activation of Notch signaling (Schweisguth, 2015). The differential activation of Notch relies on the unequal partitioning of the two cell fate determinants Neuralized (Neur) and Numb in the anterior SOP daughter cell (Le Borgne and Schweisguth, 2003; Rhyu et al., 1994). Neur promotes the endocytosis of Delta, one of the Notch ligands (Le Borgne and Schweisguth, 2003), while Numb inhibits the recycling of Notch and its cofactor Sanpodo (Spdo) towards the plasma membrane to instead promote their targeting towards late endosomal compartments (Cotton et al., 2013; Couturier et al., 2013; Johnson et al., 2016; Upadhyay et al., 2013). Consequently, the anterior cell adopts the pIIb identity while Notch is selectively activated in the posterior cell that adopts the pIIa fate. Combination of live-imaging, FRAP experiments using NiGFP and photo-tracking of photoconverted NimMaple3 revealed that proteolytic activation of Notch occurs during SOP cytokinesis and that a specific pool of Notch receptors located basal to the midbody is the main contributor to the signaling in the pIIa cell (Trylinski et al., 2017). These data imply a polarized trafficking of Notch, Delta and Spdo towards this specific subcellular location during cytokinesis.

We previously reported that the clathrin adaptor complex AP-1 regulates the polarized sorting of Notch and Spdo from the *trans*-Golgi network (TGN) and the recycling endosomes (RE) towards the plasma membrane (Benhra et al., 2011). Loss of AP-1 causes stabilization of Notch and Spdo at the adherens junctions following SOP division, a phenotype associated with a mild *Notch* gain-of-function phenotype (GOF). Unequally partitioned Numb controls the endosomal sorting of Notch/Spdo after asymmetric division and prevents their recycling to the plasma membrane (Couturier et al., 2013). This recycling event relies on AP-1 activity (Cotton et al., 2013). We also reported that Stratum (Strat), a chaperone regulating Rab8 recruitment, controls the exit from the Golgi apparatus and the basolateral targeting of Notch, Delta and Spdo (Bellec et al., 2018). As for AP-1, loss of Strat leads to an enrichment in Notch and Spdo at the apical pole of SOP daughter cells associated with a mild *Notch* GOF phenotype. As AP-1 and Strat/Rab8 both regulate Notch and Spdo trafficking to the basolateral plasma membrane, a possible interpretation of our data is that AP-1 and Strat act in the same transport pathway and therefore are simply fine-tune regulators of Notch-Delta trafficking and activation. An alternative explanation could be that AP-1 and Strat function in two parallel pathways (**Fig.1A-A’**) to ensure proper basal localization of the Notch receptor. In this scenario, loss of one of the two components could be at least in part compensated by the other. A prediction of this second hypothesis is that the concomitant loss of AP-1 and Strat would exhibit a stronger phenotype.

**Figure 1:**
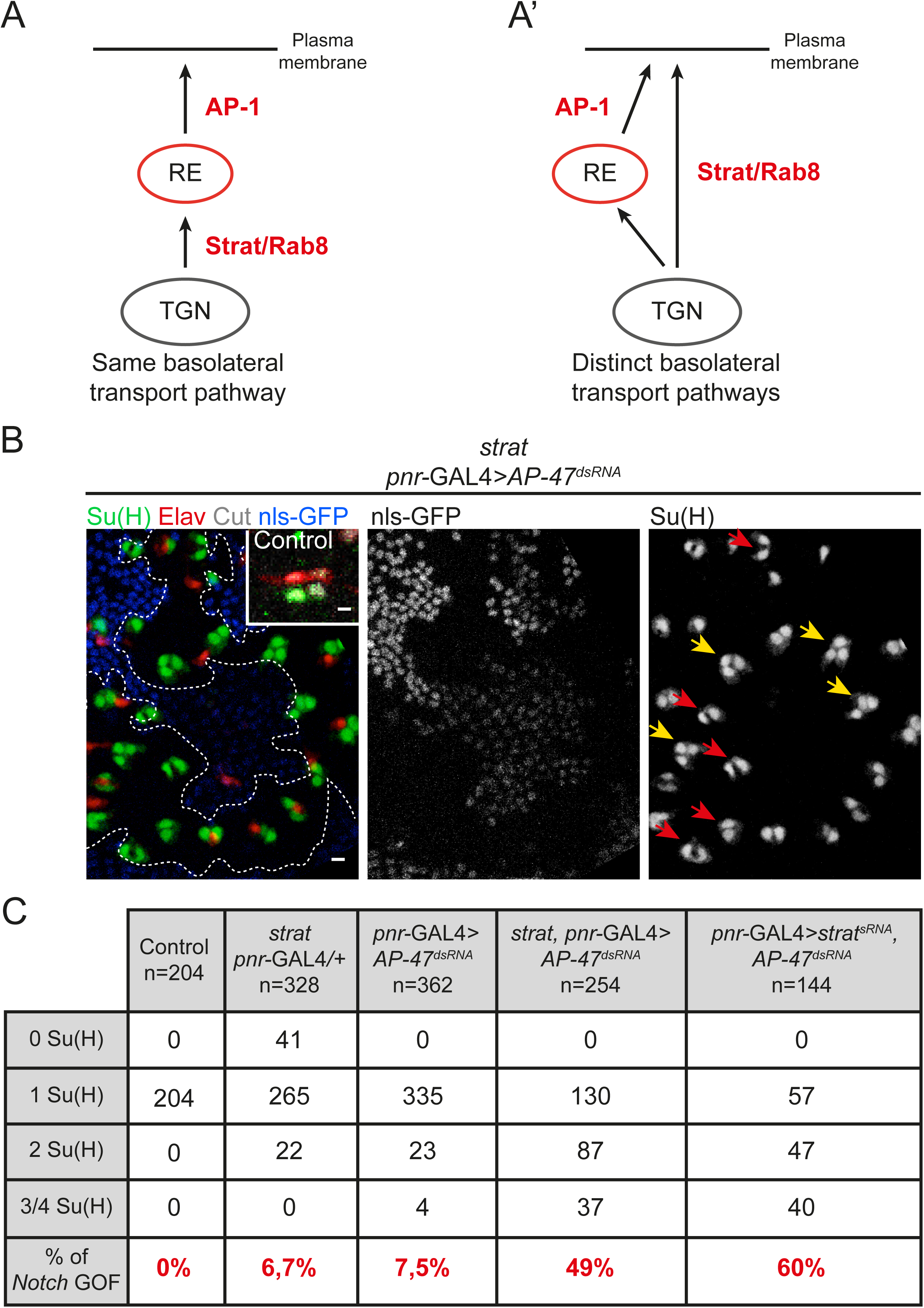
Loss of Strat and AP-1 causes *Notch* gain-of-function phenotype within SO lineage. **A-A’.** Schematic representations of the involvement of AP-1 and Strat in the same (**A**) or in distinct (**A’**) basolateral transport pathway. **B.** Projection of confocal sections of a pupal notum at 24h APF in a wild-type organ or in *strat* SOP expressing *pnr*-GAL4>*AP-47*^*dsRNA*^. *strat* mutant cells were identified by the absence of the nuclear marker nls-GFP (grey). Sockets were identified with Su(H) (anti-Su(H), green) and neurons with Elav (anti-Elav, red). In the wild-type organ, cells composing the SO are identified with Cut (anti-Cut, blue). Scale bar is 5 μm. Yellow and red arrows show transformed SOs containing 3 to 4 Su(H) positive cells or 2 Su(H) positive cells without neuron, respectively. **C.** Percentage of transformed organs at 24h APF in wild-type (n=204), in *strat, pnr*-GAL4> + (n=328), in *pnr*-GAL4>*AP-47*^*dsRNA*^ (n=362), in *strat, pnr*-GAL4>*AP-47*^*dsRNA*^ (n=254) and in *pnr*-GAL4>*strat*^*dsRNA*^, *AP-47*^*dsRNA*^ (n=144).

In this study, we investigated the consequences of simultaneous disruption of Strat and AP-1 function. We report that concomitant impairment of Strat and AP-1 does neither impact the overall apico-basal polarity of epithelial cells, nor the unequal partitioning of Numb and Neur at SOP cytokinesis. However, it does result in increased amounts of Notch, Spdo and Delta at the apical pole of the SOP daughter cells while most of Notch and Spdo, normally localized basally at the pIIa-pIIb interface (Trylinski et al., 2017), appear to be lost. This phenotype is associated with a pIIb-to-pIIa cell fate transformation that is much more penetrant than that of the single *ap-1* or *strat* mutants alone. Photoconversion and spatio-temporal monitoring of NimMaple3 in the context of simultaneous impairment of AP-1 and Strat indicate that this fate conversion may be a consequence of aberrant Notch localization. Under simultaneous loss of AP-1 and Strat, Notch receptors localized in excess at the apical pole appear to be able to translocate to nuclei of both SOP daughter cells. This could explain the *Notch* GOF phenotype as Notch activation may now occur in both SOP daughter cells. We propose a model according which AP-1 and Strat control two parallel transport routes that both contribute to the polarized transport of Notch and Spdo to the basal pIIa-pIIb interface to ensure binary cell fate acquisition at SOP cytokinesis.

## Results

### Simultaneous loss of AP-1 and Strat causes a penetrant *Notch* GOF phenotype

To test if AP-1 and Strat function in the same pathway to regulate the activation of Notch signaling pathway, we induced clones of cells homozygote mutant for a null mutation of *stra*t generated by CRISPR (Bellec et al., 2018) in which the µ subunit of the AP-1 complex (also known as AP-47) was silenced using previously characterized tools (Benhra et al., 2011). Silencing of *AP-47* is hereafter referred to as loss of AP-1. If Strat and AP-1 act in two different basolateral pathways, we expect the *Notch* GOF phenotype observed in absence of the two proteins to be much more penetrant. In the wild-type sensory organ (SO), the pIIb cell divides two times to generate the internal cells among which there is one Elav positive neuron. The pIIa cell divides once to generate the external cells among which there is one socket cell identified by Suppressor of Hairless (Su(H)). Therefore Su(H) can be used as a read-out of cell fate transformations to monitor the effect of individual or simultaneous impairment of Strat and AP-1 (**Fig.1**). In agreement with previous studies, mild *Notch* GOF phenotypes revealed by an excess of Su(H) positive socket cells were observed in 6,7% and 7,5% of SO mutant for *strat* or depleted of AP-1, respectively (Bellec et al., 2018; Benhra et al., 2011). However simultaneous loss of AP-1 and Strat led to a much stronger *Notch* GOF phenotype (49% of transformed SOs, **Fig.1B-C**). Among them, 34% of SOs had at least two Su(H) positive cells with no Elav positive cells indicating that these SOs were already transformed at the two-cell stage (**Fig.1C**). These results were confirmed using a different method, i.e. silencing both AP1 and Strat using dsRNA (**Fig.1C**). The strong enhancement of the *Notch* GOF phenotypes in this lineage analysis indicates that AP-1 and Strat act in two distinct and complementary pathways to regulate Notch activation.

### Cell polarity remodeling and unequal partitioning of cell fate determinants upon loss of AP-1 and Strat

As Strat and AP-1 regulate basolateral trafficking, we first monitored the effect of loss of these two regulators on the distribution of several cell polarity markers. We noticed that, as in *AP-1* mutants, the cuticle is thinner and less pigmented than in controls (**Fig.S1A-B’**). Despite these cuticle defects, the localization of the junctional markers DE-Cadherin, Par3 and Coracle is unaffected by the loss of AP-1 and Strat, suggesting that cell polarity is not altered (**Fig.S1C-D**). However, time-lapse imaging revealed that Par3, instead of being enriched at the posterior pole of SOP prior to mitosis (Bellaiche et al., 2001; Besson et al., 2015), distributes uniformly at the apical cortex of SOP upon loss of Strat and AP-1 at this stage of the cell cycle (orange arrows, **Fig.S1D**). This defect in planar cell polarity (PCP) regulation is likely caused by the loss of AP-1 as reported in the *Drosophila* wing (Carvajal-Gonzalez et al., 2012). Despite this defect, during mitosis, Par3 localizes normally at the posterior cortex upon loss of AP-1 and Strat, as it is the case in control SOP (**Fig. S2A and S2C**). As Par3 is required for the asymmetric localization of Numb and Neur during mitosis (Bellaïche et al., 2001; Langevin et al., 2005), we anticipated that their localization would not be affected in *strat* mutant SOPs depleted of AP-1. Indeed, we found that Numb and Neur localize asymmetrically during prometaphase in absence of Strat and AP-1 (**Fig.2A-B**). Live-imaging of Numb::GFP^crispr^ (Bellec et al., 2018) revealed that Numb is unequally partitioned in the anterior SOP daughter cell, in a similar manner as in the control situation (**Fig.2C-D**). We conclude that the *Notch* GOF phenotype is unlikely caused by defective Numb or Neur partitioning during SOP division. These data raise the possibility that the defect originates later, perhaps in the course of cytokinesis.

**Figure 2:**
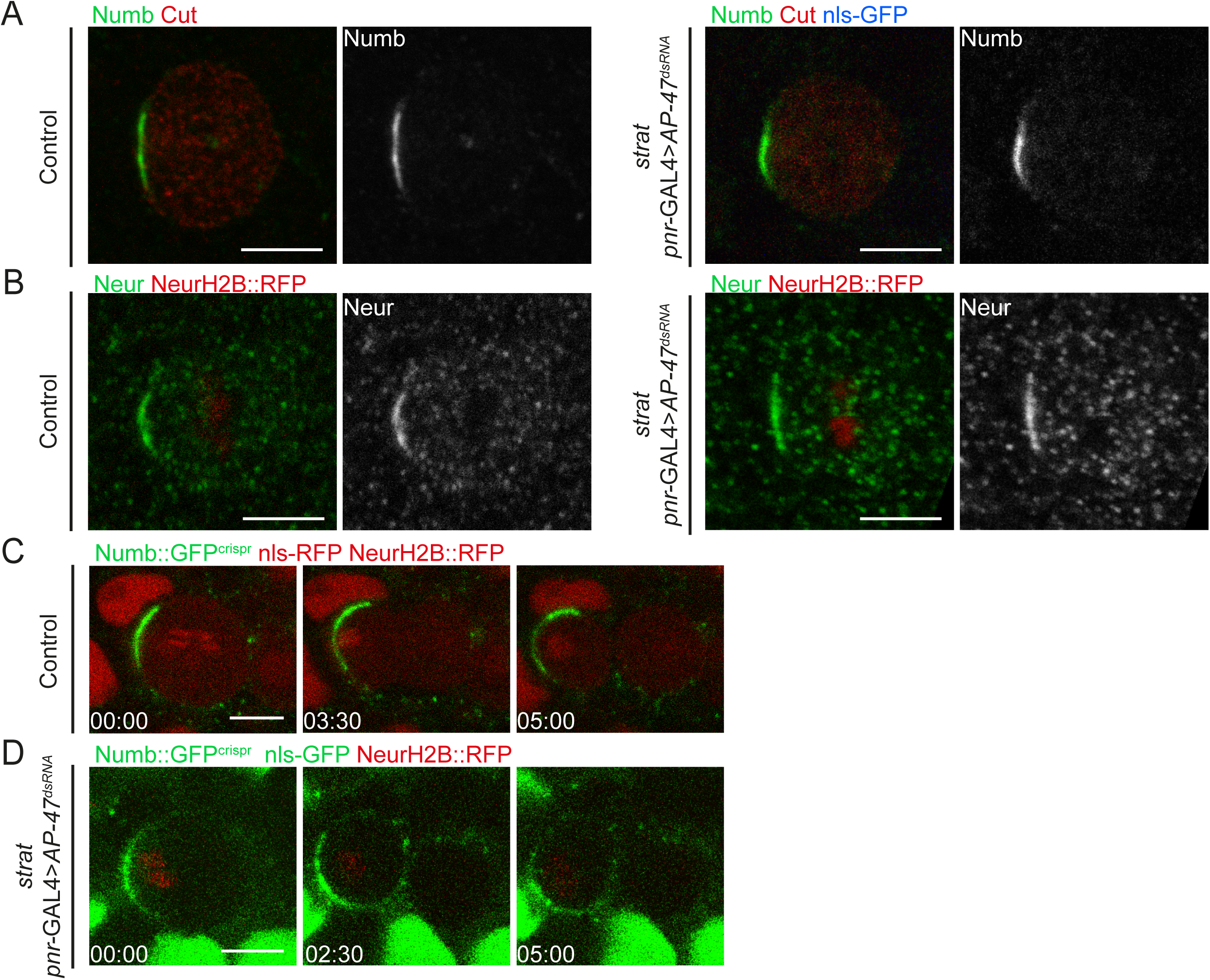
Loss of Strat and AP-1 does not affect the localization of Numb. **A.** Localization of Numb (anti-Numb, green) in wild-type (n=22 of prophase/prometaphase and dividing SOPs) and in *strat* SOP expressing *pnr*-GAL4>*AP-47*^*dsRNA*^ (n=16 of prophase/prometaphase and dividing SOPs). SOPs and SOPs daughter cells were identified with Cut (anti-Cut, red). **B.** Localization of Neur (anti-Neur, green) in wild-type (n=14 of prophase/prometaphase and dividing SOPs) and in *strat* SOP expressing *pnr*-GAL4>*AP-47*^*dsRNA*^ (n=7 of prophase/prometaphase and dividing SOPs). SOPs and SOPs daughter cells were identified with Histone2B::RFP expressed under the *neur* promoter (red). Similar phenotype was observed in the *pnr*-GAL4>*strat*^*dsRNA*^, *AP-47*^*dsRNA*^ (data not shown, n=8 of prophase/prometaphase and dividing SOPs). **C-D.** Time-lapse imaging of Numb::GFP^crispr^ (green) in dividing wild-type SOP expressing Histone2B::RFP under the *neur* promoter (red; **C**, n=10) and in dividing *strat* SOP expressing Histone2B::RFP under the *neur* promoter (red) and *pnr*-GAL4>*AP-47*^*dsRNA*^ (**D**, n=15). Time is minute:second and the time 00:00 corresponds to the SOP anaphase onset. Scale bar is 5 μm.

During cytokinesis, Par3 localizes at the apical cortex and is enriched at the posterior pole of pIIa cells, as well as at the pIIa-pIIb apical interface (**Fig.S2A**, t=21 min). In addition, we found that, in the wild-type situation, Par3 localizes in clusters at the lateral pIIa-pIIb interface (**Fig.S2A**). These Par3 clusters are also positive for Notch bearing a GFP tag in its intracellular domain (**Fig.S2A**; Houssin and Le Borgne, manuscript in preparation). Strikingly, we found that these lateral Par3/Notch clusters are absent or barely detectable upon loss of Strat and AP-1 activities (**Fig.S2C**). Instead, upon loss of Strat and AP-1, Par3/Notch clusters now reside within the apical plane of the SOP daughter cells (white arrows, **Fig.S2C**). Collectively, these data suggest that a defect in the localization of Notch could be causative to the *Notch* GOF phenotype observed upon loss of Strat and AP-1.

### Notch is enriched at the apical pIIa-pIIb interface upon loss of AP-1 and Strat

To further investigate the effect of AP-1 and Strat on the apico-basal distribution of Notch receptors, we monitored the dynamics of Notch, with a GFP tag in the Notch intracellular domain (Notch::GFP^crispr^ (Bellec et al, 2018), hereafter called NiGFP) throughout the asymmetric division of the SOP, identified by the nuclear marker Histone2B::RFP expressed under a minimal *neur* promoter (H2B::RFP, **Fig.3**). Previously, elegant work from F. Schweisguth laboratory identified two pools of Notch at the pIIa-pIIb interface, apical and basal to the midbody respectively, and provides the compelling evidence that only the subset of receptors located basal to the midbody contributes to signaling (Trylinski et al., 2017). In the control situation, we confirmed the presence of two pools of NiGFP along the apical-basal pIIa-pIIb interface (**Fig.3A-A’ and S2A**). NiGFP is transiently detected at the apical pIIa-pIIb interface at ∼6-9 min after anaphase onset with a signal intensity peaking at 15-20 min prior to progressively disappear at ∼30 min (**Fig.3A-B, S2A-B and Movie S1**). Basal to the midbody, NiGFP localizes in lateral clusters, which appear ∼6-9 min following the anaphase onset and persist longer than the apical pool of NiGFP (up to 45 min after the anaphase onset; **Fig.3A-B’ and Fig.S2A**). While NiGFP colocalizes with Par3 in these lateral clusters in wild-type (**Fig.3A-A’, 3B’ and S2A**), upon loss of Strat and AP-1, the NiGFP-Par3 lateral clusters were barely detectable (**Fig.3B’-C’ and S2C**). Instead, we found that NiGFP appears in higher amounts at the apical pIIa-pIIb interface cells ∼6-9 minutes following the anaphase onset and persists there for longer periods of time compared to the control (at least 78 min post-anaphase versus ∼30 min in the control; **Fig.3B-C’, S2C and Movie S2**). In addition to accumulating at the apical interface, NiGFP also localizes in intracellular compartments in the apical plane, some of which are positive for Par3 (white arrows, **Fig.3C-C’, S2C and Movie S2**). Since lateral clusters from which Notch is activated (Trylinski et al., 2017) are barely detectable upon loss of Strat and AP-1, our results prompt the hypothesis that the *Notch* GOF phenotype originates from the ectopic apically enriched pool of NiGFP.

**Figure 3:**
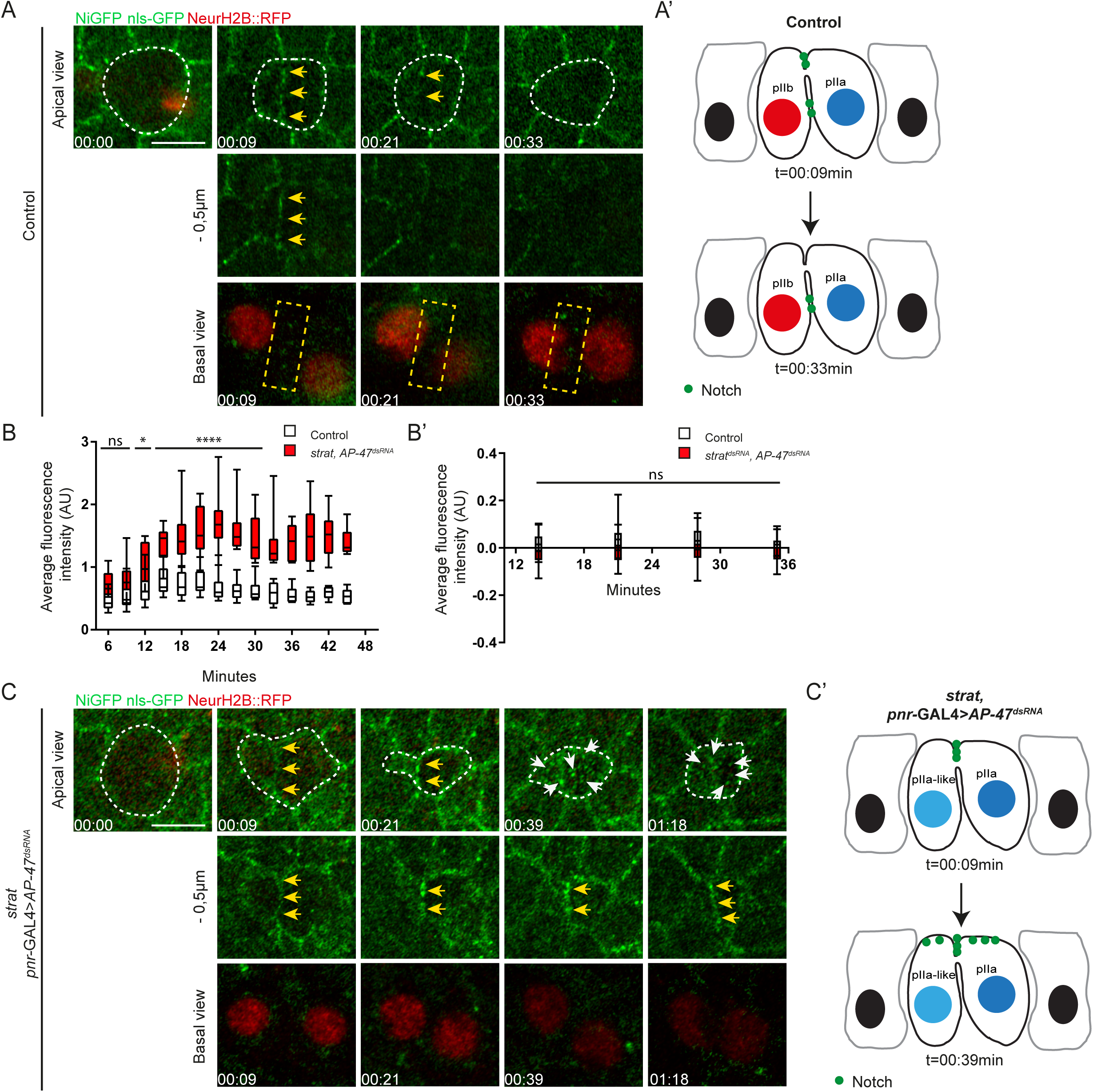
Notch is enriched at the apical pIIa-pIIb interface in the absence of Strat and AP-1. **A.** Time-lapse imaging of NiGFP (green) and Histone2B::RFP expressed under the *neur* promoter (red) in dividing wild-type SOP (n=10). **A’.** Schematic representation of Notch localization at t=9min and t=33min in wild-type SOP daughter cells. **B.** Quantification of the fluorescence intensity of NiGFP at the apical pIIa-pIIb interface of wild-type SOP daughter cells (white) and *strat* SOP daughter cells expressing *pnr*-GAL4>*AP-47*^*dsRNA*^ (red). Boxes extend from the 25^th^ to 75^th^ percentiles and the line in boxes represents the median. The whiskers go down to the smallest value and up to the largest. The statistics were carried out from t=0 to 30min and n=10 for both conditions (ns≥0.05; \**P*<0.05 and \*\*\*\**P*<0.0001). Statistics were carried out until 30min as most of wild-type SOPs movies stops at 30min. **B’.** Quantification of the fluorescence intensity of NiGFP at the basal pIIa-pIIb interface of wild-type SOP daughter cells (white, n=11) and SOP daughter cells expressing *pnr*-GAL4>*strat*^*dsRNA*^, *AP-47*^*dsRNA*^ (red, n=22). Boxes extend from the 25^th^ to 75^th^ percentiles and the line in boxes represents the median. The whiskers go down to the smallest value and up to the largest (ns≥0.05). **C.** Time-lapse imaging of NiGFP (green) and Histone2B::RFP expressed under the *neur* promoter (red) in dividing *strat* SOP expressing *pnr*-GAL4>*AP-47*^*dsRNA*^ (n=10/12). **C’.** Schematic representation of Notch localization at t=9min and t=39min in *strat* SOP daughter cells expressing *pnr*-GAL4>*AP-47*^*dsRNA*^. Dashed white lines highlight SOP and SOP daughter cells. Yellow arrows point to the enrichment of NiGFP at the apical interface between SOP daughter cells and white arrows point to apical compartments positive for NiGFP. Dashed yellow rectangles highlight compartments positive for NiGFP at the basolateral interface between wild-type SOP daughter cells. Basal views are a maximum projection of three confocal slices (Sum projection). Time is in hour:minute and the time 00:00 corresponds to the SOP anaphase onset. Scale bar is 5 μm.

### Spdo, Neur and Delta are distributed with Notch at the apical pIIa-pIIb interface upon loss of Strat and AP-1

Previously, *Notch* phenotypes observed in *AP-1* or *strat* mutant were associated with defects in Spdo and Delta localization. This prompted us to study the localization of Spdo and Delta on fixed specimens. As previously described, in the control situation, Spdo is faintly detected at the apical pole of SOP daughter cells and localizes predominantly in endosomes in the pIIb cell. In the pIIa cell, Spdo distributes not only in endosomes (yellow arrows, **Fig.4A**) but also at the basolateral plasma membrane (red arrows, **Fig.4A**; (Cotton et al., 2013; Couturier et al., 2013; Hutterer and Knoblich, 2005; Langevin et al., 2005). In contrast upon loss of Strat and AP-1, Spdo is still detected in dotted intracellular structures in SOP daughter cells but is no longer detected at the basolateral pIIa-pIIb interface (**Fig.4A**). Instead, Spdo is enriched at the apical plasma membrane as well as at the apical pIIa-pIIb interface (n=12/14, white arrows, **Fig.4A-B**).

**Figure 4:**
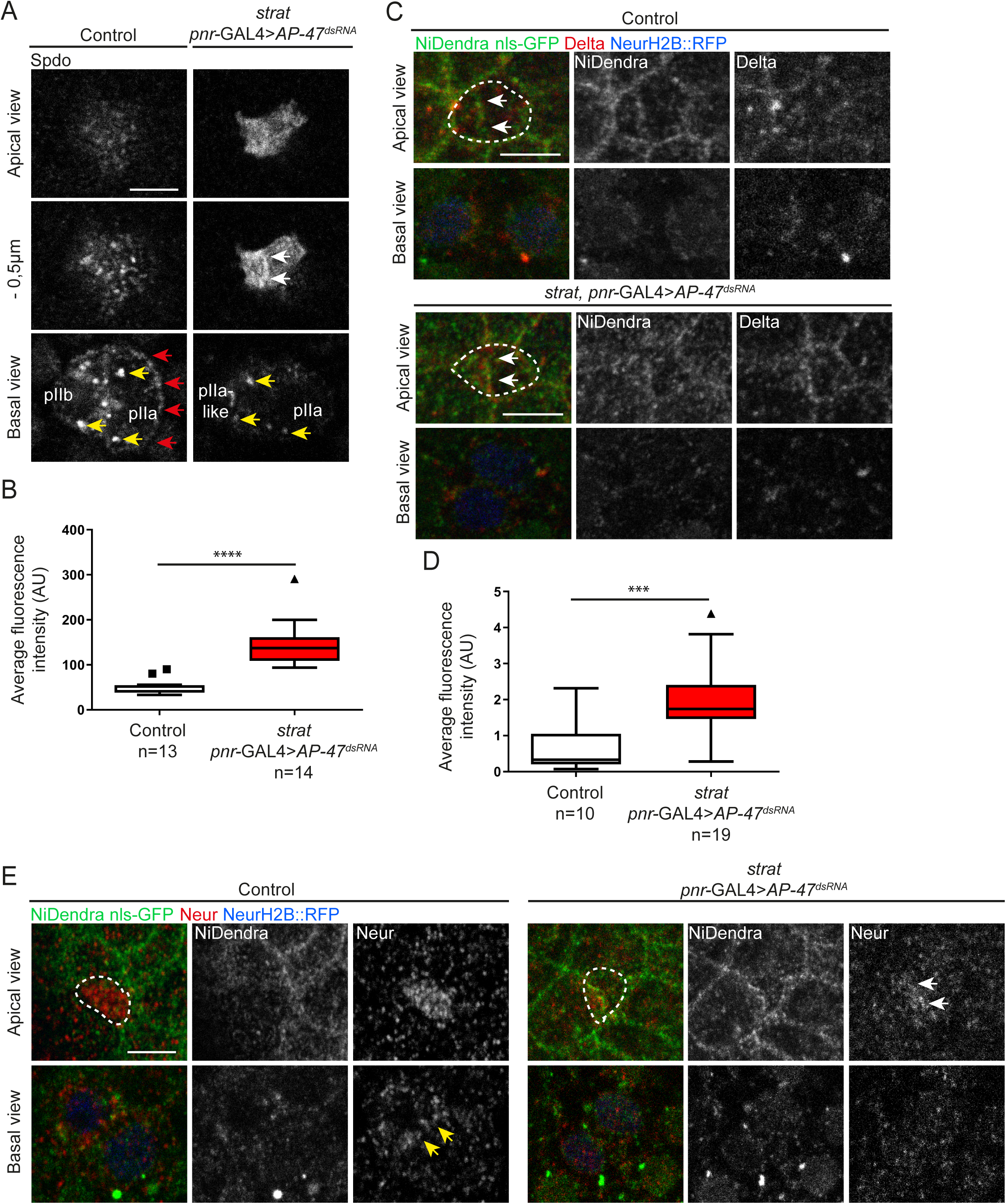
Spdo, Neur and Delta are enriched at the apical pIIa-pIIb interface in absence of Strat and AP-1. **A.** Localization of Spdo (anti-Spdo) in wild-type SOP daughter cells and in *strat* SOP daughter cells expressing *pnr*-GAL4>*AP-47*^*dsRNA*^. White arrows point to the enrichment of Spdo at the apical interface between SOP daughter cells, red arrows point the enrichment of Spdo at the basolateral plasma membrane and yellow arrows point to endosomes positive for Spdo. **B.** Quantification of the fluorescence intensity of Spdo at the apical pole of wild-type and *strat* SOP daughter cells expressing *pnr*-GAL4>*AP-47*^*dsRNA*^ (\*\*\*\**P*<0.0001). Boxes extend from the 25^th^ to 75^th^ percentiles and the line in boxes represents the median. The squares and the triangle show maximum values. **C.** Localization of NiDendra (anti-Dendra, green), Histone2B::RFP expressed under the *neur* promoter (blue) and Delta (anti-Delta, red) in wild-type SOP daughter cells and in *strat* SOP daughter cells expressing *pnr*-GAL4>*AP-47*^*dsRNA*^. White arrows point to the enrichment of Notch and Delta at the apical interface between SOP daughter cells. A projection (Maximum intensity) of the three most apical planes is shown for the apical view. **D.** Quantification of the fluorescence intensity of Delta at the apical interface of wild-type and *strat* SOP daughter cells expressing *pnr*-GAL4>*AP-47*^*dsRNA*^ (\*\*\**P*<0.001). Boxes extend from the 25^th^ to 75^th^ percentiles and the line in boxes represents the median. The triangle shows the maximum value. **E.** Localization of Neur (anti-Neur, green) and Histone2B::RFP expressed under the *neur* promoter (red) in wild-type SOP daughter cells (n=10) and in *strat* SOP daughter cells expressing *pnr*-GAL4>*AP-47*^*dsRNA*^ (n=3). The same phenotype is observed in the *pnr*-GAL4>*strat*^*dsRNA*^, *AP-47*^*dsRNA*^ (data not shown, n=9). Yellow arrows point to the enrichment of Neur at the basolateral interface between SOP daughter cells and white arrows point to the enrichment of Neur at the apical interface between SOP daughter cells. Dashed white lines highlight SOP and SOP daughter cells. Scale bar is 5 μm.

Notch activation requires Neur-mediated Delta endocytosis. In the wild-type situation, Delta and Neur localize at the lateral pIIa-pIIb interface (Trylinski et al, 2017; **Fig.4C-E**). While Neur is uniformly distributed at the apical pole in cytokinesis (Trylinski et al, 2017; **Fig.4E**), Delta is barely detected at the apical pIIa-pIIb interface (**Fig.4C-D**). The loss of Strat and AP-1 does not appear to significantly alter the location of the Neur at the pIIa-pIIb interface (**Fig.4E**) but it results in an increase in the number of SOPs with Delta labelling at the apical pIIa-pIIb interface (60% of SOs against 44% in the control situation, **Fig.4C-D**). Therefore, loss of Strat and AP-1 causes abnormally high levels of Notch/Delta and Spdo at the apical pIIa-pIIb interface at the expense of their localization to the basolateral interface. These observations raised the question of whether Delta-mediated Notch activation can take place at the apical interface upon loss of Strat and AP-1. We tested the dynamics of the ectopic apical Notch pool using photobleaching. The apical pool of NiGFP in absence of Strat and AP-1 appears to be constantly replenished and rapidly recovered (yellow arrows, **Fig.S3A**). We noticed that 120 seconds after photobleaching, the fluorescence recovery reaches its maximum in control and upon loss of Strat and AP-1. However, the mobile fraction is around 67% +/- 30% upon loss of Strat and AP-1 compared to 31% +/- 15% in a control situation (**Fig.S3B**) with a t1/2 of around 40 seconds +/- 7 seconds and a t1/2 of 37 seconds +/- 11 seconds, respectively (**Fig.SC**). The high recovery rate of the apical pool of NiGFP raises the question of its destiny and is in agreement with Delta-mediated Notch activation that can take place at the apical interface upon loss of Strat and AP-1.

### The apical pool of Notch contributes to the *Notch* GOF phenotype caused by the loss of Strat and AP-1

To further investigate the ability of the apical pool of Notch to be cleaved and transferred into the nucleus, we photoconverted Notch receptors present at the membrane and measured the presence of photoconverted Notch in the nuclei of daughter cells using NimMaple3 (Trylinski et al., 2017). We first validated our ability to photoconvert nuclear NimMaple3 in the wild-type SOP daughter cells and in SOP daughter cells deprived of Strat and AP-1, 30 minutes after the anaphase onset (**Fig.5A-B**). The plasma membrane of SOPs and daughter cells were identified using GAP43::IRFP670 (thereafter referred to as GAP43::IR) expressed under a minimal *neur* promoter (**Fig.5A’**). As reported, in a control situation, higher levels of photoconverted nuclear Notch were found in the pIIa cell compared to the pIIb cell, thus reflecting the differential activation of the Notch signaling (Trylinski et al., 2017, **Fig.5B’**). On the contrary, in absence of Strat and AP-1, the amount of photoconverted nuclear Notch is similar between the two daughter cells, in agreement with the *Notch* GOF phenotype described (**Fig.5B’**). Once photoconverted, the nuclear signal rapidly decreased over time (t1/2= 5 min), in agreement with the known instability of nuclear NICD (**Fig.S4D-D’**, (Gupta-Rossi et al., 2001; Jarriault et al., 1995)).

**Figure 5:**
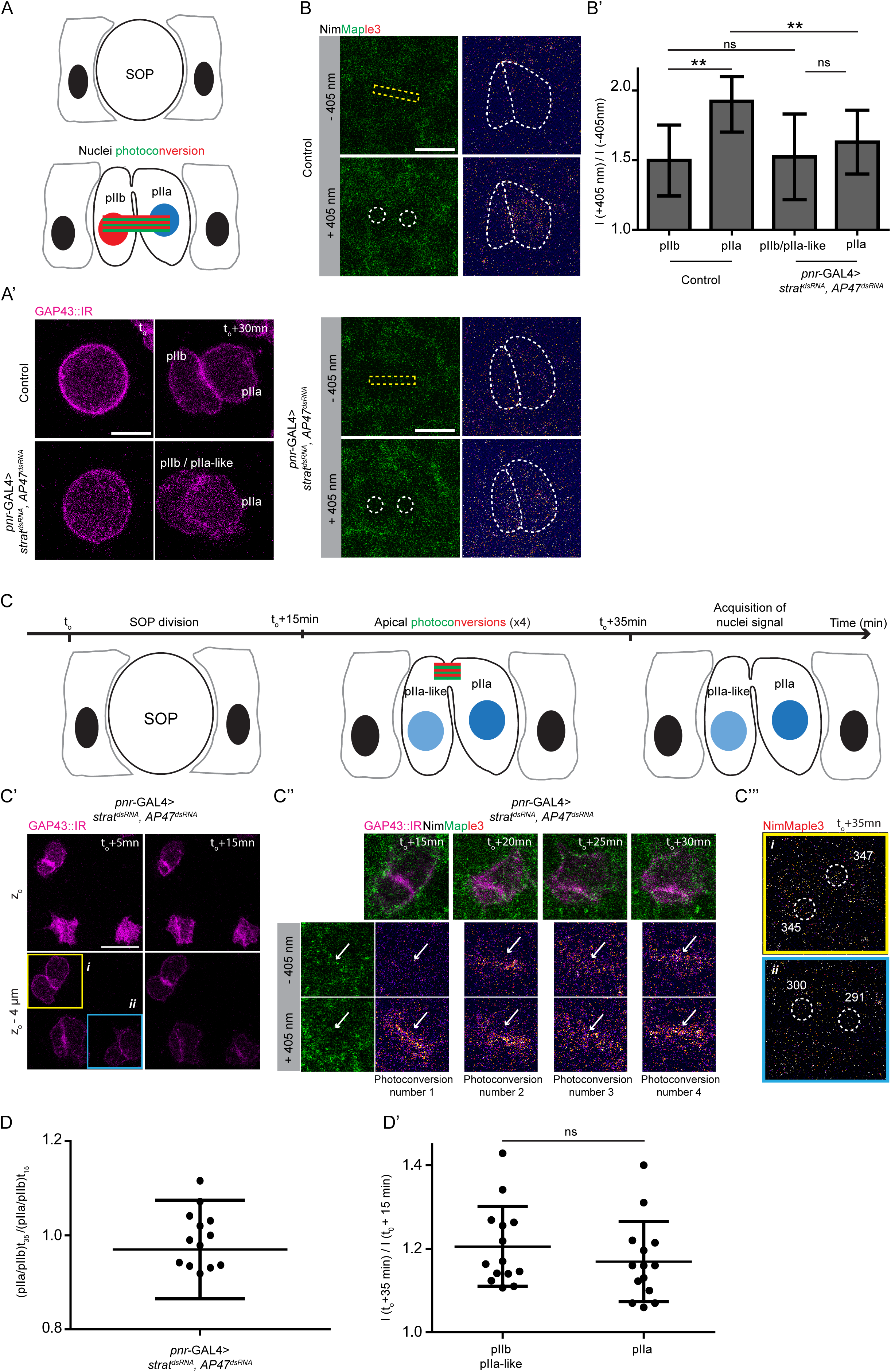
Photoconverted apical Notch is detected in SOP daughter cells nuclei in absence of Strat and AP-1. **A-A’.** Schematic representations and time-lapse imaging of wild-type and *pnr*-GAL4>*strat*^*dsRNA*^, *AP-47*^*dsRNA*^ SOP daughter cells expressing GAP43::IR. Green and red rectangles represent the photoconversion area. **B.** Photoconversion at t_o_ + 30 minutes of SOP daughter cells nuclei in wild-type and *pnr*-GAL4>*strat*^*dsRNA*^, *AP-47*^*dsRNA*^. Dashed yellow rectangles represent the photoconverted ROI, dashed white circles represent the area where nuclei signal has been measured and dashed white lines highlight SOP daughter cells. The red signal of NimMaple3 is measured before and after photoconversion. **B’.** Plot of photoconverted nuclear signal in pIIb and pIIa cells in wild-type (n=13, 3 pupae) and *pnr*-GAL4>*strat*^*dsRNA*^, *AP-47*^*dsRNA*^ (n=12, 4 pupae). **C-C’’’.** Photoconversion of apical NimMaple3 and nuclei signal measurement in *pnr*-GAL4>*strat*^*dsRNA*^, *AP-47*^*dsRNA*^ SOP daughter cells. **C.** Schematic representation of apical photoconversion assays in *pnr*-GAL4>*strat*^*dsRNA*^, *AP-47*^*dsRNA*^ SOP daughter cells. Apical photoconversions were performed at 15, 20, 25 and 30 minutes after anaphase transition. Before photoconversion, z-stacks were performed to localize the apical interface using GAP43::IR. A z-stack was then acquired 35 minutes after the anaphase onset to quantify nuclear NimMaple3. **C’.** Time-lapse imaging of *pnr*-GAL4>*strat*^*dsRNA*^, *AP-47*^*dsRNA*^ SOPs cells expressing GAP43::IR before apical photoconversions. Yellow square highlights the SOP where apical Notch is photoconverted and the blue square highlights the SOP where Notch is not photoconverted and serves as reference. **C’’.** Photoconversion of apical NimMaple3 at t_o_ + 15, 20, 25 and 30 minutes in *pnr*-GAL4>*strat*^*dsRNA*^, *AP-47*^*dsRNA*^ SOPs cells expressing GAP43::IR (n=14, 4 pupae). White arrows show Notch present at the apical pIIa-pIIb interface. **C’’’.** Measurements of nuclei signal (nuclei are delimited by dashed white circles) in photoconverted cell (upper panel, yellow) and non-photoconverted cell (lower panel, blue). pIIb or pIIa-like are localized at the left of the apical interface and pIIa are localized at the right of the apical interface. **D.** pIIa/pIIb ratio values of photoconverted nuclear NimMaple3 at t_o_ + 30 minutes upon apical NimMaple3 photoconversion in *pnr*-GAL4>*strat*^*dsRNA*^, *AP-47*^*dsRNA*^ (n=14, 4 pupae). **D’.** Plot of photoconverted nuclear signal in pIIb/pIIa-like and pIIa cells in *pnr*-GAL4>*strat*^*dsRNA*^, *AP-47*^*dsRNA*^ (n=14, 4 pupae). Scale bar is 5 µm. t_o_ represents the SOP anaphase onset. In **B’** and **D-D’**, data are represented as mean +/- SD. (ns≥0.05 and \*\**P*<0.01). Because GAP43::IR is excluded from the nuclei, nuclei were defined using an area within cells where the GAP43::IR signal background is the lowest.

We then tested the precision of NimMaple3 photoconversion in controls. Photoconversion at the level of the apical pIIa-pIIb interface remained restricted to the apical plane as no undesired photoconversion occurred deeper in the cell (**Fig.S4A-A’’**). In contrast, photoconversion within a ROI encompassing the lateral pIIa-pIIb plasma membrane interface lead to unwanted photoconversion in the adjacent nuclei (**Fig.S4B-B’’**) and at the apical level over a diameter of about 3-4 cells (**Fig.S4C-C’**). Therefore, the inability to ensure precise photoconversion on the lateral interface prevents testing by photoconversion whether Notch is activated from the apical, the lateral or from both pools of receptors in controls. This technical caveat is, however, not a concern upon loss of Strat and AP-1 as NimMaple3 signal (or NiGFP) is barely detected at the lateral interface (**Fig.3B’-C’ and Fig.S2C**). When we photoconverted Notch at apical pIIa-pIIb interface in this context, we found that Notch can enter in both nuclei that result from the division (**Fig.5C-C’’**). The levels of photoconverted Notch measured in both nuclei (**Fig.5C”’**, cell *i* highlighted in yellow on panel **5C’**) are further higher than those found in the nuclei of a neighbouring SOP that divides in the same time, but that was not exposed to photoconversion (**Fig.5C’’’**, cell *ii* highlighted in blue on panel **5C’**). On average, the contribution of the apical pool of Notch when Strat and AP-1 are simultaneously impaired seems identical between the two nuclei as demonstrated by the ratio of photoconverted nuclear signal pIIa/pIIa-like close to 1 and the absence of significant differences in the amount of photoconverted nuclear signal between the two cells (**Fig.5D-D’**). Taken together, these data demonstrate that, in absence of Strat and AP-1, the apical enrichment of Notch at the interface is source of activated receptors present in nuclei of both daughter cells therefore causing *Notch* GOF phenotype.

## Discussion

### Strat and AP-1 act redundantly in the basolateral transport of Notch signaling components in SOPs

Previous work showed that Strat/Rab8 control the transport of Notch, Delta and Spdo from the TGN to the basolateral pole (Bellec et al., 2018) and that AP-1 controls the targeting of Notch and Spdo from the TGN to the basolateral membrane via the recycling endosomes (Benhra et al., 2011; Cotton et al., 2013). Here, we report that upon concomitant loss of Strat and AP-1, Notch, Delta and Spdo are no longer detected in clusters at the lateral SOP daughter cells interface and are instead enriched at the apical interface (**Fig.6**; Bellec et al., 2018; Benhra et al., 2011; Cotton et al., 2013). We found that AP-1 and Strat act redundantly to control the basolateral targeting of Notch, Delta and Spdo in SOP daughter cells explaining the partial compensation of one by the other. These results are in agreement with previous studies demonstrating the involvement of Rab8, whose activity and localization is controlled by Strat in *Drosophila* (Bellec et al., 2018; Devergne et al., 2017), in the basolateral sorting of proteins (Ang et al., 2003; Henry and Sheff, 2008; Huber et al., 1993). However the role of Rab8is not restricted to the basolateral transport as the loss of one or the other has also been previously associated to defects in apical targeting and secretion (Castillon et al., 2018; Gillard et al., 2015; Holloway et al., 2013; Nakajo et al., 2016; Norgate et al., 2006; Sato et al., 2014; Sato et al., 2007; Zhang et al., 2012). In the same way, the role of AP-1 is not restricted to the basolateral transport. In our study, we described a thinner and less pigmented cuticle in absence of Strat and AP-1. These phenotypes could be caused by a defective AP-1 dependent transport of the Menkes Copper transporter ATP7a that regulates cuticle pigmentation (Holloway et al., 2013; Norgate et al., 2006) and/or by a reduction in apical secretion of cuticle components that may rely on AP-1 function as does the apical glue granule secretion in salivary glands (Burgess et al., 2011). These apparent contradictory results might be explained by a distinct Rab8- or AP-1-mediated sorting depending on the cargo to be transported or on the tissue. In addition, sorting might occur at the level of RE. If a protein that does not normally transit via RE is found to be present in RE in a mutant background, it may be sorted and transported to an unusual apical location by default.

**Figure 6:**
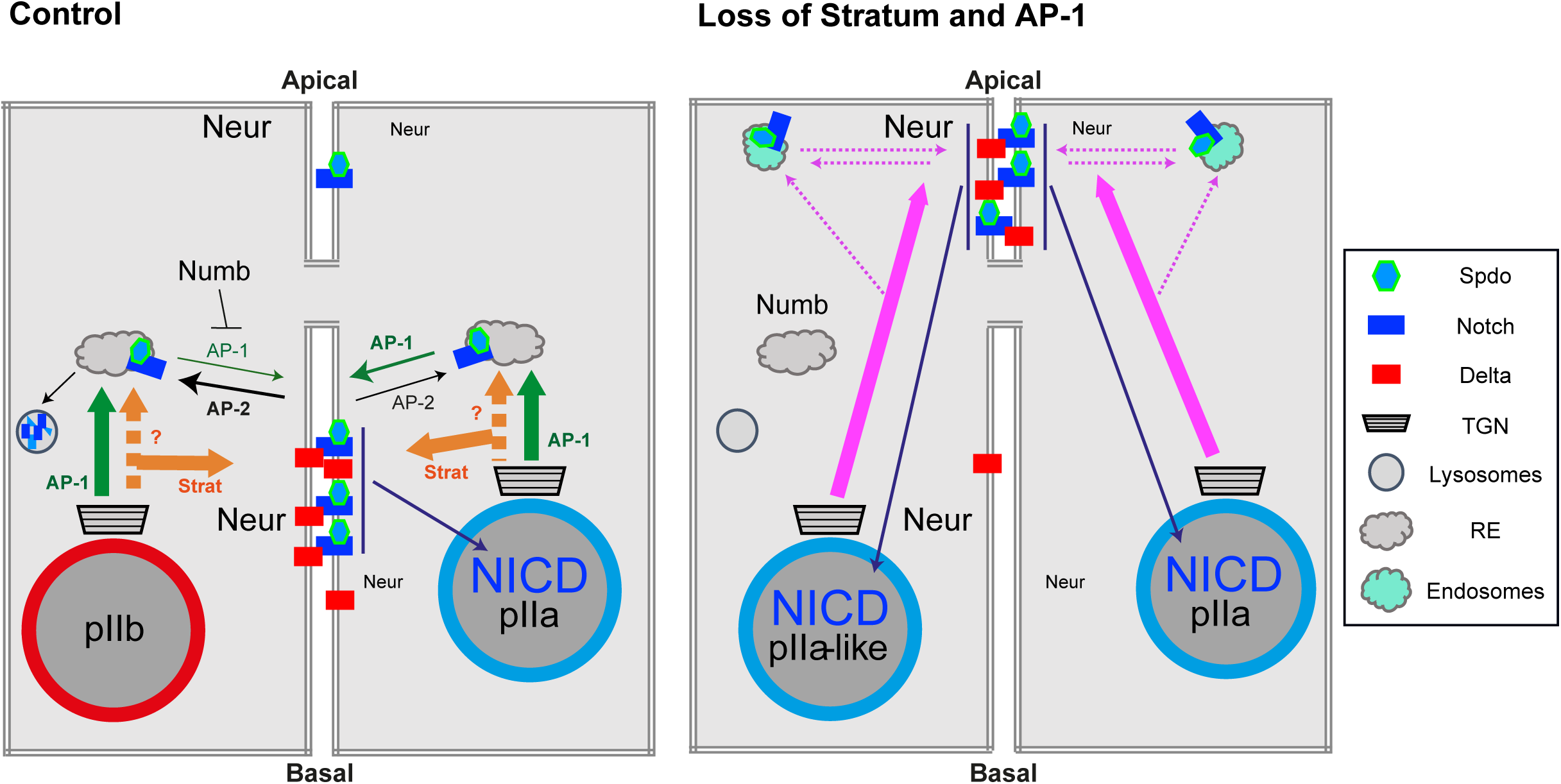
Model for the roles of Strat and AP-1 in the transport of Notch signaling components. Schematic representation of Strat and AP-1 function in the transport of Notch, Delta and Spdo in wild-type and in absence of Strat and AP-1. Nuclei of cells are represented with a blue circle (Notch positive) or in red (Notch negative). AP-2, which is asymmetrically enriched in pIIb (Berdnik et al., 2002), promotes the endocytosis of Notch/Spdo (black arrows), while AP-1 regulates their recycling to basolateral membrane (green arrows), a step negatively regulated by Numb in pIIb cells. The traffic of Notch, Delta and Spdo that is dependent on Strat and AP-1 is represented in orange or in green arrows, respectively. In absence of Strat and AP-1, the mistrafficking of Notch, Delta and Spdo is represented by pink arrows. Notch and Spdo are also found in apical dotted compartment, and their possible routing to and from there is depicted by pink dashed arrow (TGN: for the *trans*-Golgi network and RE: recycling endosomes).

### Notch signaling activation can take place at the apical interface of the SOP daughter cells

How can Notch be activated in the absence of lateral clusters, the main contributors of intra-lineage Notch activation (Trylinski et al., 2017), in absence of Strat and AP-1? Our work indicates that Notch activation can take place at the apical pIIa-pIIb interface. Indeed, photo-tracking of photoconverted NimMaple3 in the *strat* mutant background deprived of AP-1 revealed that the apical pool of Notch is an active pool of Notch in terms of signaling, contributing to the nuclear NICD signal in both daughter cells. However, we cannot exclude the possibility that a few, hard to detect, lateral clusters could also contribute to Notch activation upon loss of AP-1 and Strat. In the same way, we cannot exclude that Notch activation can take place from the Notch positive compartments observed at the apical pole of SOP daughter cells upon loss of Strat and AP-1.

The fact that Sec15, a subunit of the exocyst complex, and Arp2/3-WASp (Actin-related protein 2/3-Wiskott-Aldrich syndrome protein) are involved in apical trafficking of Spdo and Delta, respectively, and are required for Notch-Delta signaling is consistent with our proposal of apical activation of Notch (Jafar-Nejad et al., 2005; Rajan et al., 2009). However, the weak contribution of the apical pool of receptors to the Notch activation described in the control situation (Trylinski et al., 2017) suggests that apical activation can only occur when a given threshold of Notch, Delta and Spdo is reached, such as in *strat* SOs depleted of AP-1. In line with previous reports (Trylinski et al., 2017), our study further illustrates how the apico-basal targeting of Notch, Spdo and Delta along the pIIa-pIIb interface limits signaling between SOP daughter cells at the exit of mitosis. The question of whether this mechanism is conserved awaits further investigation, but given the key role played by Notch in the acquisition of cell fate in vertebrates, it is conceivable that the regulation described in this study may be general.

### Loss of Strat and AP-1 causes Notch activation in both SOP daughter cells

As reported in this study, the loss of Strat and AP-1 leads to a *Notch* GOF despite the unequal partitioning of Numb. This phenotype can be explained by the lack of AP-1 that prevents Numb to repress the Notch-Spdo recycling normally driving endosomal degradation of Notch-Spdo (Cotton et al., 2013; Couturier et al., 2013; Johnson et al., 2016). Thus, Numb is not able to repress Notch activation upon loss of AP-1 and Strat, thereby unable to trigger symmetry breaking in Notch signaling at SOP cytokinesis.

Proteolytic activation of Notch requires the binding of the ligand and its subsequent Neur-mediated endocytosis. While Neur is unequally localized during SOP division (Le Borgne and Schweisguth, 2003), Neur::GFP was shown to localize in both SOP daughter cell apical cortex and to be enriched at the apical pIIa-pIIb interface following SOP division (Trylinski et al., 2017). As Delta and Neur localizes at the apical SOP daughter cell interface upon loss of Strat and AP-1 (this work), we propose that Neur can promote endocytosis of Delta in both daughter cells from the apical interface, thus leading to the bidirectional Notch proteolytic activation. In agreement with this proposal according which Neur might be active in the two SOP daughter cells, it is interesting to note that in *numb* mutant cells, despite the unequal inheritance of Neur, Delta-dependent Notch activation takes place in both daughter cells, as observed here upon loss of Strat and AP-1.

## Acknowledgements

We thank the Bloomington Stock Center, the Vienna Drosophila RNAi Center and the National Institute of Genetics Fly Stock Center for providing fly stocks. We also thank S. Dutertre and X. Pinson from the Microscopy Rennes Imaging Center-BIOSIT (France). The monoclonal antibodies against Elav, Cut, Cora and DE-Cad were obtained from the Developmental Studies Hybridoma Bank, generated under the auspices of the National Institute of Child Health and Human Development, and maintained by the University of Iowa Department of Biological Sciences. We thank Dr. J. Januschke, University of Dundee for sharing the Par3::Scarlet *Drosophila* line prior to publication. We thank Dr. J. Januschke and the members of R.L.B.’s laboratory for helpful discussions and critical reading of the manuscript.

## Competing interests

The authors declare no competing or financial interests.

## Author contributions

Conceptualization: K.B., R.L.B.; Methodology: K.B., M.P., I.G., R.L.B.; Formal analysis: K.B., M.P., R.L.B.; Investigation: K.B., M.P., R.L.B.; Resources: K.B., I.G., R.L.B.; Writing - original draft: K.B.; Writing - review & editing: K.B., M.P., R.L.B.; Visualization: K.B., M.P., R.L.B.; Supervision: R.L.B.; Project administration: R.L.B.; Funding acquisition: K.B., R.L.B.

## Funding

This work was supported in part by the ARED programme from the Région Bretagne/Agence Nationale de la Recherche (ANR-16-CE13-004-01), the Fondation pour la Recherche Médicale (FDT20170436864 to K.B.), La Ligue contre le Cancer-Equipe Labellisée (R.L.B.) and the Association Nationale de la Recherche et de la Technologie programme PRC Vie, santé et bien-être CytoSIGN (ANR-16-CE13-004-01 to R.L.B.).

## Materials and Methods

### *Drosophila* stocks and genetics

*Drosophila melanogaster* stocks were maintained and crossed at 25°C. Mitotic clones were induced using the FLP-FRT technique using the *hs*-FLP and by heat shocking (2×60 min at 37°C) at second and early third instar larvae. *pnr*-GAL4 was used to drive the expression of the *AP-47*^*dsRNA*^ and the *strat*^*dsRNA*^. The following stocks were used in this study:

**Figure 1B:** *y, hs*-FLP (/*w*); *Ubi-GFP nls*, FRT40A/*Ubi-GFP nls*, FRT40A and *y, w, hs*-FLP (/*w*^*-*^); *Ubi-GFP nls*, FRT40A/*strat*, FRT40A; *pnr-*GAL4/*AP-47*^*dsRNA*^ (stock #24017 from Vienna Drosophila Resource Center, *w*^*1118*^; P(GD14206)v24017/TM3)

**Figure 1C:** *y, hs*-FLP (/*w*); *Ubi-GFP nls*, FRT40A/*Ubi-GFP nls*, FRT40A and *y, hs*-FLP (/*w*); *Ubi-GFP nls*, FRT40A/*strat*, FRT40A and *y, w, hs*-FLP (/*w*); *Ubi-GFP nls*, FRT40A/*strat*, FRT40A; *pnr-*GAL4/*AP-47*^*dsRNA*^ and *y, w, NimMaple3*/Y; *strat*^*dsRNA*^ /+ ; *pnr-*GAL4, *neur*-GAP43::IRFP670/*AP-47*^*dsRNA*^

**Figure 2A:** *Cad::GFP*/+; *pnr-*GAL4/+; +/CyO::CFP; *pnr*-GAL4/+ and *y, w, hs*-FLP (/*w*); *Ubi-GFP nls*, FRT40A/*strat*, FRT40A; *pnr-*GAL4/*AP-47*^*dsRNA*^

**Figure 2B:** *y, w, hs*-FLP/ *NiDendra, neur-H2B*::RFP; *Ubi-GFP nls*, FRT40A/FRT40A; *pnr-* GAL4/+ and *y, w, hs*-FLP/*NiDendra, neur-H2B*::RFP; *Ubi-GFP nls*, FRT40A/*strat*, FRT40A; *pnr-*GAL4/*AP-47*^*dsRNA*^

**Figure 2C:** *y, hs*-FLP (*w*); *Ubi-RFP nls*, FRT40A/*Numb*::*GFP*^crispr^, FRT40A

**Figure 2D:** *w, hs*-FLP *neur-H2B*::RFP/*w*; *Ubi-GFP nls*, FRT40A/ *strat, Numb*::*GFP*^crispr^, FRT40A; *pnr-*GAL4/ *AP-47*^*dsRNA*^

**Figure 3:** *y, w, hs*-FLP/*NiGFP, neur-H2B*::RFP; *Ubi-GFP nls*, FRT40A/FRT40A; *pnr-*GAL4/+ and *y, w, hs*-FLP/*NiGFP, neur-H2B*::RFP; *Ubi-GFP nls*, FRT40A/*strat*, FRT40A; *pnr-*GAL4/ *AP-47*^*dsRNA*^ and *y, w, NiGFP, Par3::Scarlet*; *+/+*; *pnr-*GAL4/*+* and *y, w, NiGFP, Par3::Scarlet*;*strat* ^*dsRNA*^ */+*; *pnr-*GAL4/*AP-47*^*dsRNA*^

**Figure 4A-B:** *W1118* and *y, w, hs*-FLP (/*w*); *Ubi-GFP nls*, FRT40A/*strat*, FRT40A; *pnr-*GAL4/*AP-47*^*dsRNA*^

**Figure 4C-D:** *y, w, hs*-FLP/ *NiDendra, neur-H2B*::RFP; *Ubi-GFP nls*, FRT40A/FRT40A; *pnr-* GAL4/+ and *y, w, hs*-FLP/*NiDendra, neur-H2B*::RFP; *Ubi-GFP nls*, FRT40A/*strat*, FRT40A; *pnr-*GAL4/*AP-47*^*dsRNA*^

**Figure 4E:** *y, w, hs*-FLP/*NiDendra, neur-H2B*::RFP; *Ubi-GFP nls*, FRT40A/FRT40A; *pnr-* GAL4/+ and *y, w, hs*-FLP/*NiDendra, neur-H2B*::RFP; *Ubi-GFP nls*, FRT40A/*strat*, FRT40A; *pnr-*GAL4/*AP-47*^*dsRNA*^ and *y, w, NiGFP, Par3::Scarlet*; *strat*^*dsRNA*^/+; *pnr-*GAL4/*AP-47*^*dsRNA*^

**Figure 5:** *y, w, NimMaple3*/Y; *strat* ^*dsRNA*^ /+ ; *pnr-*GAL4, *neur*-GAP43::IRFP670/*AP-47* ^*dsRNA*^

**Figure S1A-A’:** *NiDendra, neur-H2B*::RFP; +/+

**Figure S1B-B’:** *y, w, hs*-FLP/*NiDendra, neur-H2B*::RFP; *Ubi-GFP nls*, FRT40A/*strat*, FRT40A; *pnr-*GAL4/*AP-47*^*dsRNA*^

**Figure S1C:** *y, w, hs*-FLP (/*w*^*-*^); *Ubi-GFP nls*, FRT40A/*strat*, FRT40A; *pnr-*GAL4/*AP-47*^*dsRNA*^

**Figure S1D:** *y, w, NiGFP, Par3::Scarlet*; *+/+*; *pnr-*GAL4/*+* and *y, w, NiGFP, Par3::Scarlet*; *strat*^*dsRNA*^/+; *pnr-*GAL4/*AP-47*^*dsRNA*^

**Figure S2:** *y, w, NiGFP, Par3::Scarlet*; *+/+*; *pnr-*GAL4/*+* and *y, w, NiGFP, Par3::Scarlet*;*strat* ^*dsRNA*^ */+*; *pnr-*GAL4/*AP-47*^*dsRNA*^

**Figure S3A:** *y, w, NiGFP, Par3::Scarlet*; *strat*^*dsRNA*^/+; *pnr-*GAL4/*AP-47*^*dsRNA*^

**Figure S3B-C:** *y, w, NiGFP5, neur-H2B*::RFP (control) and *y, w, NiGFP, Par3::Scarlet*; *strat*^*dsRNA*^/+; *pnr-*GAL4/*AP-47*^*dsRNA*^

**Figure S4A-C:** *y, w, NimMaple3*/Y; + /+ *pnr-*GAL4, *neur*-GAP43::IRFP670/+

**Figure S4D-D’:** *y, w, NimMaple3*/Y; *strat* ^*dsRNA*^/+ ; *pnr-*GAL4, *neur*-GAP43::IRFP670/*AP-47* ^*dsRNA*^

**Movie S1:** *y, w, hs*-FLP/ *NiGFP, neur-H2B*::RFP; *Ubi-GFP nls*, FRT40A/FRT40A; *pnr-*GAL4/+

**Movie S2:** *y, w, hs*-FLP/*NiGFP, neur-H2B*::RFP; *Ubi-GFP nls*, FRT40A/*strat*, FRT40A; *pnr-*GAL4/ *AP-47*^*dsRNA*^

### Immunofluorescence and antibodies

Pupae were aged for 16.5 to 18.5 h after puparium formation (APF) for SOPs and SOPs daughter cells analysis and were aged for 24 h to 28 h APF for lineage analysis. Pupae were dissected in 1x phosphate-buffered saline (1x PBS) and then fixed for 15 min in 4% paraformaldehyde at room temperature. Dissection and staining conditions were essentially as previously described (Le Borgne and Schweisguth, 2003). Primary antibodies used were rat anti-Elav (7E10, Developmental Studies Hybridoma Bank (DSHB), 1:200), goat anti-Su(H) (sc15813, Santa Cruz, 1:500), mouse anti-Cut (2B10, DSHB, 1:500), rabbit anti-Neur ((Lai et al., 2001), 1:1000), goat anti-Numb (SC23579, Santa Cruz, 1:200), rabbit anti-Spdo (a kind gift from J. Skeath, 1:2000) (O’Connor-Giles and Skeath, 2003), mouse anti-DECD (C594.9B, DSHB, 1:200), mouse anti-Cora (C615.16, DSHB, 1:500), rat anti-DE-Cad (DCAD2, DSHB, 1:500) and rabbit anti-Dendra (Antibodies-online.com, ABIN361314, 1:1000). Cy2-, Cy3- and Cy5-coupled secondary antibodies (1:400) were from Jackson’s Laboratories.

### Generation of NiDendra

The NiDendra construct was generated using the CRISPR/Cas9 method as previously described (Gratz et al., 2013a; Gratz et al., 2013b). As for the NiGFP construct, the following gRNAs were used: 5′-AACTTGAA|TGGATTGAACCC GGG-3′ and 5′-CGAACTGG|AGGGTTCTCCTGTTG-3′ to introduce Cas9 cuts in exon 6 (Bellec et al., 2018). The Dendra and the 3xP3-DsRed cassette flanked by GVG linkers and by loxP, respectively, were introduced at the previously described position (NiYFP4) (Couturier et al., 2012). Homology arms 1 and 2 were of 1064 bp and 1263 bp in length, respectively. Injection was performed by Bestgene in the *yw*; attP40(nos-cas9)/Cyo stock. The correct position of the Dendra and the DsRed cassette was verified by PCR and sequencing. The DsRed was then removed by crossings with *if*/Cyo, Cre, *w* stock. The insertion of Dendra tag did not alter the functionality of Notch. Anti-Dendra signal is detected in the pIIa nucleus indicating that processed NiDendra is translocated into the nucleus (data not shown).

### Imaging

Images of fixed nota were acquired with a Leica SPE confocal microscope and a Zeiss Airyscan microscope (LSM 880 with AiryScan module). Live imaging of NiGFP with Par3::Scarlet was performed with a Leica SPE confocal microscope. Live imaging of NiGFP in wild-type SOP and in *strat* SOP expressing *pnr*-GAL4>*AP-47*^*dsRNA*^ (**Fig.S2**) was performed with a Zeiss Airyscan microscope (LSM 880 with AiryScan module). All images were processed and assembled using ImageJ 1.48 and Adobe Illustrator.

### Quantification of the enrichment of Notch at the apical and basal interface

To quantify the signal of NiGFP at the apical interface, we measured the signal in a manually drawn area on sum slices of the two apical planes where the Notch signal at the apical interface is the strongest. We also measured the signal between two epithelial cells within an equivalent drawn area and on the same sum slices. Then, we calculated the following ratio: Average fluorescence intensity at the apical interface between SOP daughter cells/Average fluorescence intensity at the apical interface between epithelial cells. This ratio has been calculated for each time point. In absence of AP-1 and Strat, the manually drawn area was minimized to avoid considering the apical punctate structures positive for Notch.

To quantify the signal of NiGFP at the basal interface, we measured the signal in a manually drawn area between nuclei on sum slices of two planes. In a wild-type situation, we did the sum slices on planes where we observed basolateral clusters positive for Par3 and Notch. In absence of AP-1 and Strat and in absence of clusters, we did the sum slices on equivalent planes located between the two nuclei. In parallel, we measured the background fluorescence noise on the same sum slices, with the same manually drawn area, in the cytoplasm of epidermal cells. Then, we calculated the following ratio: Average fluorescence intensity at the basal interface between SOP daughter cells – the background fluorescence noise /Average fluorescence intensity at the apical interface between epithelial cells. This ratio has been calculated for each time point.

### Quantification of the apical enrichment of Spdon and Delta

The fluorescence intensity was calculated with ImageJ, on two-cell stages, as previously described (Bellec et al., 2018). For Spdo, the average fluorescence intensity was measured in a manually drawn area on sum slices of the two most apical planes, where Spdo is enriched. The background noise was measured in the same way and subtracted from the apical intensity value. For the Delta, the same protocol was applied: the average fluorescence intensity was measured in a manually drawn area on sum slices of the two apical planes where Delta is enriched at the interface between the two daughter cells. These values were divided by the average intensity measured at the apical interface between epidermal cells, on the same sum slices and with the same manually drawn area. The background noise was measured in the same way and subtracted from the apical intensity values.

### FRAP experiments

Photobleaching of NiGFP/NiGFP5 was performed using a Zeiss AiryScan (LSM 880 with AiryScan module) with the 488 nm laser wavelength (20 mW) at 90% of the maximal power and by using the 63x oil objective (NA 1.4). Two consecutive iterations were performed. The FRAP area was defined by the pIIa/pIIb interface. A control area corresponding to an adjacent epidermal cell interface was measured to obtain the general photobleaching of the sample over the period of acquisition. All FRAP data were analyzed using the easyFRAP software tool (https://easyfrap.vmnet.upatras.gr/?AspxAutoDetectCookieSupport=1). Half-time t1/2 and mobile fraction were then extracted with GraphPAD Prism software using a one-component equation.

### Photoconversion experiments

Photoconversion of NimMaple3 was performed using a Zeiss AiryScan (LSM 880 with AiryScan module) with a 405 nm diode (30mW) at 1.8% power (40 iterations and pixel dwell time: 1.52 µs) according to Trylinski et al., 2017.

In nuclear photoconversion assays (**Fig.5A-B**), the photoconverted ROI (dashed yellow rectangles) were defined using the GAP43::IR expressed under the *neur* promoter. Because GAP43::IR is excluded from the nuclei, nuclei were defined using an area within cells where the GAP43::IR signal background is the lowest. Nuclei measurement (dashed white circles, **Fig.5B and Fig.5C’’’**) were then performed using a circular ROI corresponding to the half of the presumptive diameter of nuclei with the center of the ROI positioned on the center of mass of nuclei.

Apical photoconversions in *pnr*-GAL4>*strat*^*dsRNA*^, *AP-47*^*dsRNA*^ SOP daughter cells were performed at 15, 20, 25 and 30 minutes after anaphase transition Before photoconversion, z-stacks were performed to localize the apical interface using GAP43::IR only first, and then with green NimMaple3. A z-stack was then acquired 35 minutes after the anaphase onset to quantify nuclear NimMaple3. No threshold has been applied.

## Statistical analysis

Statistical analyses were carried out using the GraphPad Prism 6.07 software. A two-way ANOVA was performed for the quantification of the enrichment of Notch at the apical and at the basal pIIa-pIIb interface with a multiple comparison Bonferroni test. For other quantifications, we performed a t test if the data followed a normal distribution or a Wilcoxon test or a Mann-Whitney test if the data did not follow a normal distribution. The normal distribution was tested by a Shapiro test. Statistical significances are represented as follows: not significant (ns)≥0.05; \**P*<0.05; \*\**P*<0.01; \*\*\**P*<0.001 and \*\*\*\**P*<0.0001.

